# A High-Throughput SPR-Based Array for Quantitative Profiling of Glycosaminoglycan–Protein Interactions

**DOI:** 10.64898/2026.07.02.736113

**Authors:** Holly L. Birchenough, Jonathan F. Popplewell, Douglas Dyer, Anthony J. Day, Thomas A. Jowitt

**Affiliations:** Manchester Cell-Matrix Centre, Faculty of Biology, Medicine and Health, University of Manchester, Manchester Academic Health Science Centre, Manchester, United Kingdom; Carterra Inc., Salt Lake City, Utah, USA; Lydia Becker Institute of Immunology and Inflammation, Faculty of Biology, Medicine and Health, University of Manchester, Manchester Academic Health Science Centre, Manchester United Kingdom; Geoffrey Jefferson Brain Research Centre, Manchester Academic Health Science Centre, Northern Care Alliance NHS Group, University of Manchester, Manchester Academic Health Science Centre, Manchester, United Kingdom

**Keywords:** glycosaminoglycans, GAG-protein interactions, high-throughput screening, binding assay, proteoglycan

## Abstract

Glycosaminoglycans (GAGs) are linear, negatively charged, polysaccharides that mediate a wide variety of biologically critical interactions with proteins, underpinning growth factor signalling, extracellular matrix assembly and numerous disease processes. However, GAG-protein interactions remain under characterised, in part because of the lack of high-throughput tools to systematically profile binding across the GAG interactome. In this paper we present a novel Surface Plasmon Resonance-based array methodology utilising 16 commonly sourced GAG preparations (including chondroitin sulphate (CS), dermatan sulphate (DS), heparan sulphate, heparin, hyaluronan and keratan sulphate) allowing the specificity and affinity of GAG-binding proteins to be determined. As proof of principle, we have validated the array using four established GAG-binding proteins (antithrombin III, CD44, heavy chain 1 from inter-α-inhibitor and Slit2), generating data consistent with the known binding specificities and quantifying affinities for many of the interactions. The array also reveals previously unreported GAG interactions, including Slit2 binding to CS and DS, and CD44 binding to chondroitin sulphate E.

## Introduction

Glycosaminoglycans (GAGs) are a large and diverse class of polysaccharides found in all mammalian tissues. Well known as major constituents of extracellular matrix (ECM) they are also found on the surface of all human cells (1–3). These long, linear, negatively charged polysaccharides are composed of repeating disaccharide units that, with the exception of hyaluronan (HA), can be variably modified by acetylation, epimerisation and sulphation. These modifications yield a high degree of structural heterogeneity, which in turn underpins diverse functional interactions with a variety of proteins including growth factors, chemokines, ECM components, proteases, viral proteins and more (4–8).

It is has been reported that there are over 3,500 proteins in the human “GAG-interactome” that bind one or more GAG species (9,10). Yet despite this, studies into GAG-protein interactions have been fewer compared to analogous areas such as protein-protein or protein-DNA interactions. High-throughput profiling of GAG-protein binding remains somewhat limited. This likely arises because of the complexity of GAG structures, derived from: (i) their chemical heterogeneity (chain length, epimerisation, sulphation pattern); (ii) the challenge of immobilising or presenting GAGs in a biologically relevant format, and (iii) limitations in assay throughput and quantification. The ability to systematically dissect the specificity (11) of GAG-binding proteins experimentally, as well as improving throughput whilst enabling the accurate quantification of interaction affinities would significantly advance our understanding of the field (12). This could in turn unlock new insights into biology, pathology and therapeutic targeting of GAG-protein systems (for example in cancer, fibrotic and inflammatory diseases) (13).

The complex structure of sulphated GAGs results from the numerous modifications that can be generated during biosynthesis. Referred to as the “glycocode”, these variable patterns confer structural and functional diversity (9,11,14). In this regard, GAG biosynthesis is a non-template driven process mediated by a wide range of specific enzymes (2,15), which engender GAG structures with differences in sugar type, chain length, glycosidic bond configuration and the degree and positions of sulphate groups. While HA is the exception, in that it is just composed of a repeating disaccharide (of glucuronic acid (GlcA) and N-acetyl glucosamine (GlcNAc)) without any biosynthetic modifications and exists as a free sugar not attached to a core protein, it can exhibit great heterogeneity in its molecular weight being up to ∼10 MDa in size (> 25,000 disaccharides). None-the-less it still mediates diverse biological activities through its interactions with HA-binding proteins (16,17). The sulphated GAGs (comprising chondroitin sulphate (CS) and the related dermatan sulphate (DS) formed by epimerisation of CS, heparan sulphate (HS) and its highly sulphated form, heparin, and keratan sulphate (KS)) are generally composed of ∼20-200 disaccharides and are synthesised attached to core proteins.

HS and heparin are the most structurally heterogeneous GAGs with ∼1.2 and ∼2.5 sulphates, respectively, per disaccharide (18); sulphates are principally found on the N-position of glucosamine (GlcN), replacing the N-acetyl group of GlcNAc, the 3-O and 6-O positions of GlcNS and 2-O position of iduronic acid (IdoA) after its epimerisation from GlcA. CS/DS exhibit less structural diversity with the principle isomers found in mammalian tissue having sulphation at the 4 or 6 positions of N-acetyl galactosamine (GalNAc) leading to the CS subtypes of C4S (GlcA-GalNAc4S; also known as CSA), C6S (GlcA-GalNAc6S; also known as CSC) and DS (IdoA-GalNAc4S; also called CSB) (19), sulphation can also occur on both 4 and 6 positions of GalNAc to generate the rarer CSE isoform. Sulphation of KS can occur at the 6-O position of either its galactose or GlcNAc saccharides (20).

The structural diversity of GAGs is important in mediating specific interactions and functions. For example, it has been demonstrated that differential sulphation of HS chains provides considerable selectivity in the binding of chemokines to cells, which could represent a means of regulating their localisation in tissues (21). Other examples include basic fibroblast growth factor (FGF2) which requires 6-O sulphated HS to bridge with its receptor and form its biologically active ternary complex (22,23). Antithrombin III (ATIII) requires 3-O-sulphation, in the context of a specific pentasaccharide sequence, for high affinity binding to heparin and subsequent anticoagulant activity through allosteric regulation (24). Studies have also reported the preference of Slit2 for HS sulphation sequences that contain 6-O-sulphation and N-sulphation and their subsequent effects on Slit-mediated axon guidance (25).

A variety of methods have been developed to study GAG-protein interactions (Table 1). These include low throughput techniques such as affinity chromatography (26,27) and microtitre plate-based glycosaminoglycan arrays (28,29) where, for example, various GAGs can be immobilised onto allylamine-coated surfaces. However, the data obtained are largely qualitative and, moreover, these plates are no longer commercially available.

More informative technologies include biolayer interferometry (BLI) and surface plasmon resonance (SPR). Within these methods one of the interaction partners (usually the GAG) is immobilized on a sensor surface, while the other (usually the protein) is passed over the surface (30,31). Changes in the interference pattern (BLI) or refractive index (SPR) caused by the binding of the protein to the GAG are measured in real time, providing quantitative data on the interaction kinetics and affinities. Though highly sensitive and providing quantitative data, these methods in their usual formats are not particularly high throughput. Structural techniques have also been employed for the characterisation of protein-GAG interactions. Both NMR spectroscopy and X-ray crystallography of GAG-protein complexes can produce high-resolution structures, or experimentally-inferred models, providing insights into binding modes, orientations and GAG-induced conformational changes (32–35). Though highly informative these techniques are not quantitative and by their nature low throughput.

Within recently developed ‘slide-based’ microarrays, GAG sequences or derivatives are immobilised on a slide surface and probed, e.g., with fluorescently labelled target proteins (36,37). These arrays, which generally require small sample volumes, overcome the limitations of throughput with the ability to investigate the binding of a target protein to multiple GAG variants. However, this type of method often only produces binding ‘hits’, and in some cases an estimate of *K*_D_ but not the full interaction kinetics (i.e., on and off rates that provide a more accurate determination of affinity).

A further limitation for microarrays and plate-based methods is the immobilisation strategy for the GAGs (31). Numerous methods exist for the attachment of GAGs to solid supports, both covalent and non-covalent. Non-covalent methods include immobilisation via ionic interactions of the negatively charged GAG chains with electropositive surfaces (e.g., poly-L-lysine or allylamine-coated). Covalent methods often require the chemical modification of the GAG, where for example carbodiimide chemistry (EDC/NHS) activates carboxyl groups on the GAG to form a bond with amine groups on the sensor surface or the use of glutaraldehyde as a crosslinker to react with amine groups on both the GAG and the surface. These methods of immobilisation do not closely mimic the physiological presentation of flexible/dynamic GAG chains and can occlude binding sites (38). Surface density of the immobilised GAG also needs to be optimised for accurate analysis of the protein interaction (31). This is in part to avoid steric hinderance from densely packed GAG chains blocking potential binding sites and to allow the best fitting of acquired data.

**Table 1.**
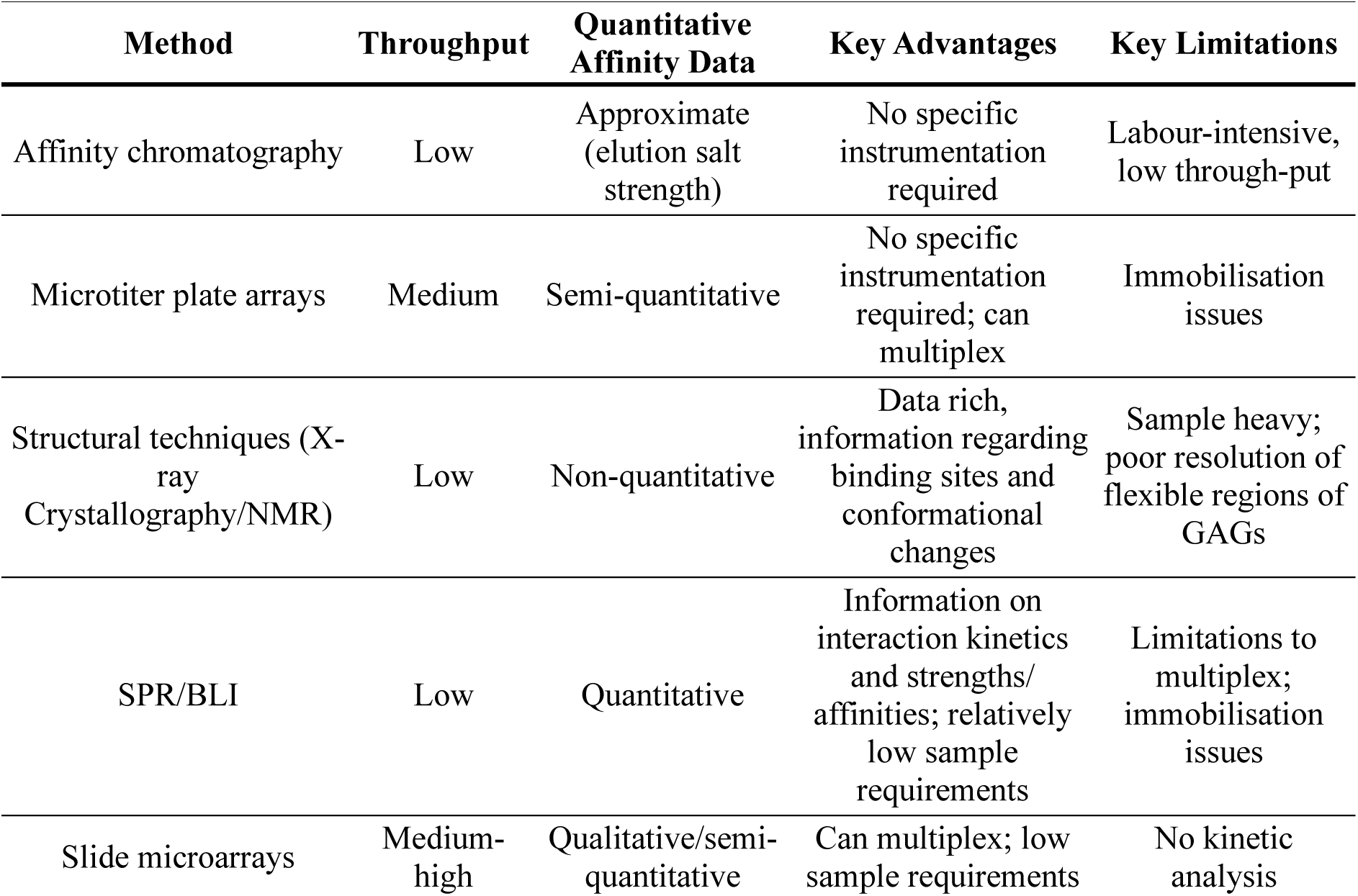
Summary of Current Methods to Analyse GAG-Protein Interactions.

| Method | Throughput | Quantitative Affinity Data | Key Advantages | Key Limitations |
| --- | --- | --- | --- | --- |
| Affinity chromatography | Low | Approximate (elution salt strength) | No specific instrumentation required | Labour-intensive, low through-put |
| Microtiter plate arrays | Medium | Semi-quantitative | No specific instrumentation required; can multiplex | Immobilisation issues |
| Structural techniques (X-ray Crystallography/NMR) | Low | Non-quantitative | Data rich, information regarding binding sites and conformational changes | Sample heavy; poor resolution of flexible regions of GAGs |
| SPR/BLI | Low | Quantitative | Information on interaction kinetics and strengths/affinities; relatively low sample requirements | Limitations to multiplex; immobilisation issues |
| Slide microarrays | Medium-high | Qualitative/semi-quantitative | Can multiplex; low sample requirements | No kinetic analysis |

To address the limitations of these GAG binding techniques there is a need for a high-throughput binding assay capable of multiplex screening of proteins against a library of GAGs, in parallel, with optimised conditions for the determination of specificity and the accurate quantification of affinity.

Here we describe a new high-throughput SPR-based method for the analysis of GAG-protein interactions utilising the Carterra LSA^XT^ instrument (Figure 1). The LSA^XT^ has a 96-multichannel fluidics system enabling up to 96 ligands to be immobilised simultaneously onto discreet spots on a sensor chip surface, which can then be repeated up to 3 times to produce a 384-spot array (Figure 1). This configuration is particularly advantageous as it allows for the immobilisation of multiple ligands at different ligand densities. A single flow channel then draws sample through a needle and passes it over all the immobilised ligands. For both the immobilisation of ligands and the flow of the sample over the chip, the solutions are oscillated back and forth, reducing the volume and thus amount of sample required (39). A high-resolution CCD camera images the entire chip surface concurrently allowing for up to 384 interactions to be analysed in parallel.

To highlight the application of this new GAG-array methodology, we have focussed here on the 4 main types of GAGs: CS/DS, HA, heparin/HS and KS (Table 2), including GAG preparations with varied chain lengths and sulphation patterns. As proof of principle, we have assayed known GAG-binding proteins, antithrombin III (ATIII), CD44, inter-α-inhibitor (IαI) heavy chain 1 (HC1) and Slit2, to illustrate the capabilities of this approach.

**Figure 1.**
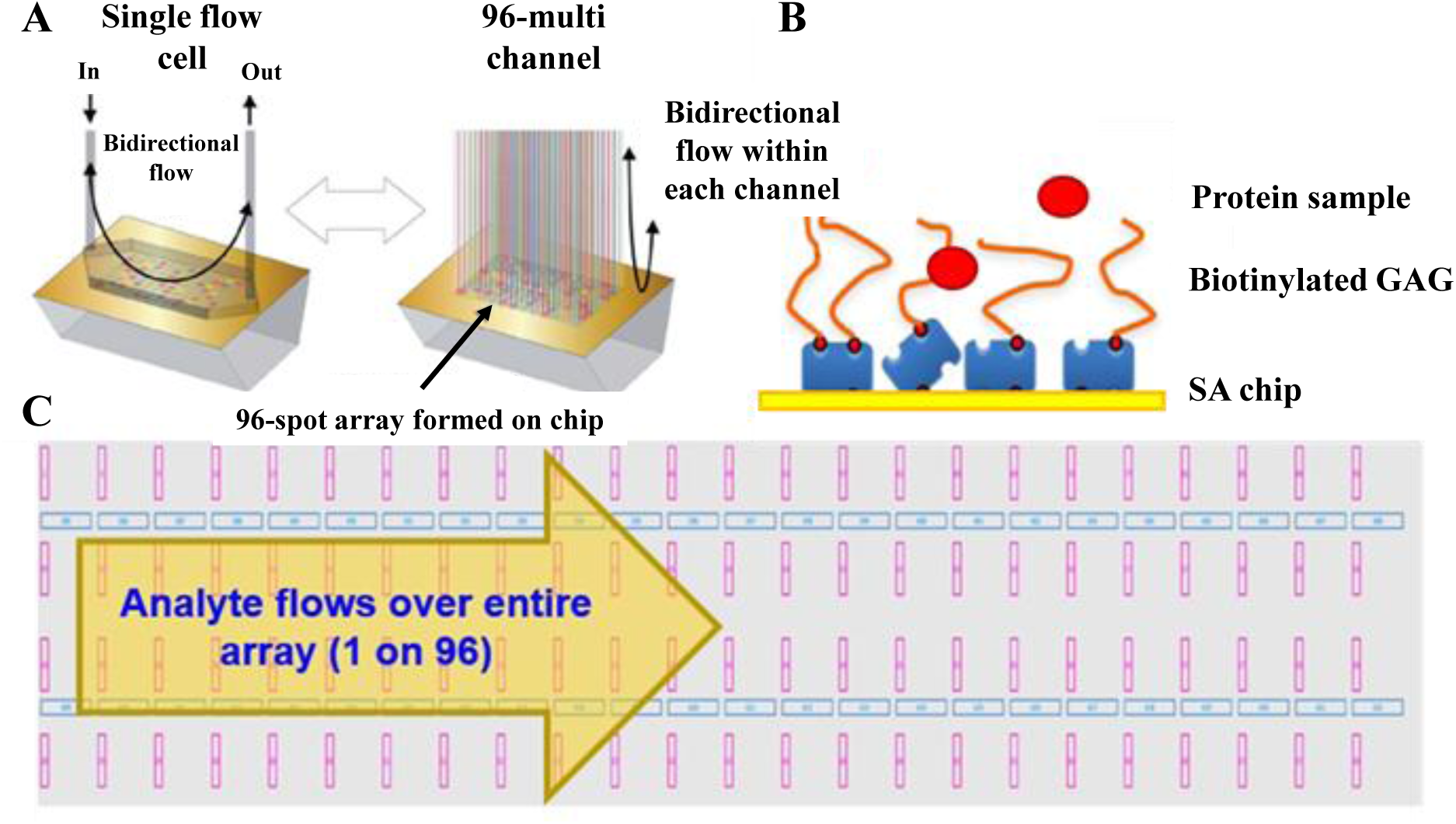
Formation of the GAG array on the Carterra LSA^XT^. (A) The Carterra LSA^XT^ operates via a single flow cell and 96-channel fluidics. (B) The multi-channel fluidics enables the formation of the 96 ‘spot’ GAG array on the sensor surface in which reducing end biotinylated GAGs (bGAG) are captured onto the streptavidin (SA) chip surface at varied capture levels. (C) The single flow cell then injects protein analyte over the entire array within a single step allowing simultaneous measurements of all GAG-protein interactions; the protein can be flowed at sequentially higher concentrations to obtain interaction parameters. Pink rectangles represent active ‘spots’ with captured GAG and blue rectangles represent reference ‘spots’ with no captured GAG.

**Table 2.** GAGs immobilised on sensor chip surface.

| GAG | Abbr. | Description | Tissue source | GAG | Abbr. | Description | Tissue source |
| --- | --- | --- | --- | --- | --- | --- | --- |
| **Heparin dp4** | dp4<sup>a</sup> | Heparin 4-mer | Porcine mucosa | **Fully desulphated heparin** | FdeS | Unfractionated heparin; all sulphate groups removed | Porcine mucosa |
| **Heparin dp8** | dp8 | Heparin 8-mer | Porcine mucosa | **Heparan sulphate** | HS | Unfractionated Heparan sulphate | Porcine mucosa |
| **Heparin dp16** | dp16 | Heparin 16-mer | Porcine mucosa | **Dermatan Sulphate** | DS | Unfractionated Dermatan sulphate | Porcine mucosa |
| **Heparin dp24** | dp24 | Heparin 24-mer | Porcine mucosa | **Keratan Sulphate** | KS | Unfractionated Keratan sulphate | Shark cartilage |
| **Unfractionated (parental) heparin** | UFH | Unfractionated heparin | Porcine mucosa | **Chondroitin Sulphate E** | CSE | GalNAc is sulphated at both the 4- and 6-positions | Shark cartilage |
| **2-O desulphated heparin** | 2OdeS | Unfractionated heparin with sulphate group removed from the C2 of IdoA | Porcine mucosa | **Chondroitin -4-Sulphate** | C4S | GalNAc is sulphated at 4-position | Bovine cartilage |
| **6-O desulphated heparin** | 6OdeS | Unfractionated heparin with sulphate group removed from the C6 of GlcN | Porcine mucosa | **Chondroitin -6-Sulphate** | C6S | GalNAc is sulphated at 6-position | Bovine cartilage |
| **N-desulphated, re-N-acetylated heparin** | NdeS | Unfractionated heparin with N-desulphation; amino group is modified by acetylation | Porcine mucosa | **Hyaluronan HA12** | HA | HA 12-mer | Bacterial |

<sup>a</sup>dp = degree of polymerisation

## Materials and Methods

### Materials

Heparin preparations and heparan sulphate (from porcine mucosa) were purchased from Iduron Ltd (Manchester, UK). Chondroitin sulphates were purchased from Sigma Aldrich (St Louis, USA). Keratan sulphate was purchased from Cosmo Bio Co Ltd (Toyko, Japan). Dermatan sulphate was provided by Barbara Mulloy (NIBSC, UK) (40). Hyaluronan 12-mer (HA12) and recombinant human CD44 HA-binding domain residues 20-169 (CD44_HABD^20-169^) were prepared as described previously (16). Recombinant human Slit2 (carrier free (CF)) was purchased from Bio-techne Ltd (UK). Recombinant human HC1 was produced according to the protocol previously described (41). Recombinant human antithrombin III (CF) was purchased from R&D systems. All other reagents were purchased from Sigma-Aldrich (St Louis, USA).

### Reducing end biotinylation of GAGs

Each of the 16 GAG preparations (Table 2) were covalently modified with biotin at their reducing end using a previously described approach (38). In brief, GAGs at 2 mg/mL in 50 mM sodium acetate were individually incubated with 20 mM aniline and 10 mM EZ-Link Alkoxyamine-PEG4-Biotin (Thermofisher) overnight at 37°C. Excess biotin reagent was removed by extensive dialysis using 7 kDa MWCO dialysis units into deionised H_2_O or by size exclusion chromatography into PBS pH 7.4 using a Superdex-75 10/30 column for dp4 and dp8 heparin.

### GAG array

Experiments were performed on a Carterra LSA^XT^ (39) using a SAHC30M chip with a running buffer of PBS, pH 7.4 with 0.05% (v/v) Tween-20 (PBST). Chips were conditioned with a 60 second injection of 40 mM NaOH. Each biotinylated GAG (bGAG) was titrated from approximately 0.375 µg/mL (dp4 and dp8) or 10 µg/mL (all other GAGs) in a 6 point, 5-fold, dilution series in PBST in a 96 deep well plate. The 6 concentrations for each bGAG (16 in total) were captured onto individual regions of interest (i.e., 96 ‘spots’) on one quadrant of the streptavidin-coated surface with a 10-minute contact time using the 96 multi-channel device.. All of the proteins analysed here were prepared in PBST with the concentration series defined in the results section. Prior to their injection onto the GAG array, an initial set of 6 PBST injections were performed as blanks. Each individual protein was then flowed over the GAG array (at up to 8 increasing concentrations) using the single needle injector with an association time of 5 minutes and a dissociation time of 15 mins without regenerations between each protein concentration. A newly captured 96-spot array was used for each individual protein, unless the previous protein sample dissociated completely (with no non-specific binding) in which case the GAG surface would be reused for the next protein under study.

### Data analysis

The data were processed using the Carterra Kinetics analysis software. Each binding curve was processed by referencing to the closest (blank) streptavidin spot, double referenced using PBST injections, and then fitted to a 1:1 Langmuir binding model with t_o_ floated, consistent with performing non regenerative kinetic analysis. Alternatively, the steady state binding model was used where appropriate. For ‘interactions’ with low/no binding responses or data that did not fit to the 1:1 model, the following quality control criteria were applied: tiles shaded grey did not meet the minimum response (10 RU) i.e., no binding, tiles shaded yellow denote residuals > 5% of R_max_ suggesting kinetic parameters generated cannot be considered a good fit to the 1:1 model, and tiles shaded blue denote that the fitted R_max_ is greater than 50% of the observed maximum response, which suggests that the K_D_ maybe higher than the highest analyte concentration and estimates maybe insufficiently constrained.

## Results

All GAGs within the array presented here were site specifically mono-biotinylated via their reducing end to allow orientated capture onto a streptavidin sensor surface. To overcome the issues of steric hinderance and analyte rebinding from high immobilisation levels, a dilution series was made for each biotinylated GAG within a 96 well plate. The 96-multichannel microfluidics system then allows the orientated capture of each GAG dilution (Figure 2). The varied immobilisation levels, seen for each of the GAGs, increase the likelihood of obtaining a suitable signal, with good on and off kinetics (i.e., for at least one coating ‘concentration’) leading to high quality data.

**Figure 2.**
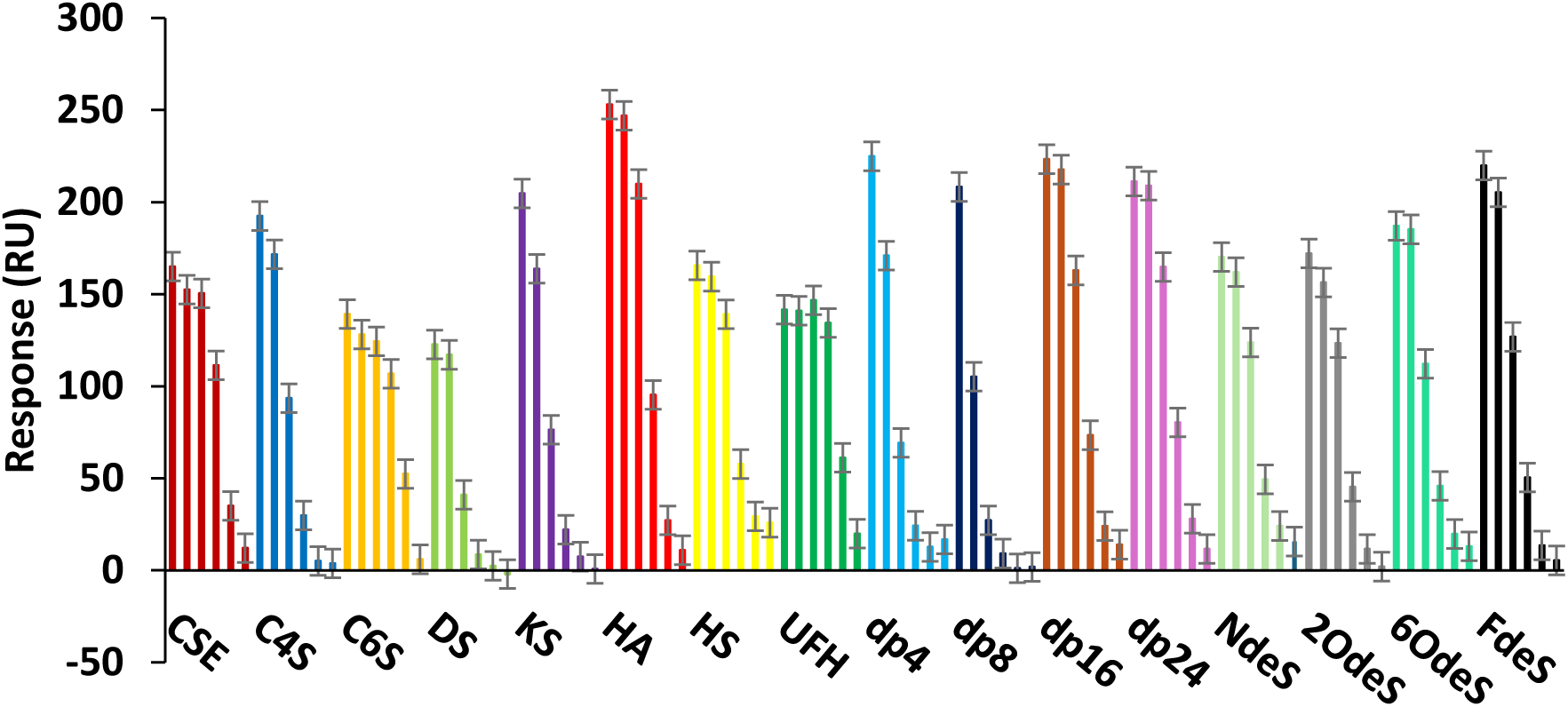
GAG capture levels on SA sensors. Six point 5-fold dilutions of reducing-end biotinylated GAGs were captured onto a single quadrant (96 spots) of streptavidin-coated sensors to create ranges of immobilisation levels. Mean GAG levels are plotted ± SD, where these are derived from six independently captured arrays.

Following capture of the GAGs onto the chip surface binding and kinetic analysis can be performed by injecting a concentration series of the potential binding partner (8 dilutions are generally used sequentially ranging from low to high concentration). Here the concentration series will vary depending on the protein; for high-quality SPR data the concentration should be at a minimum several fold above the *K*_D_ for a particular interaction and create a range of dose responses during the injection time In cases of initial screening, where the *K*_D_ (and perhaps whether the protein binds GAGs) is unknown, the maximum concentration available is recommended as the starting point for the dilution series. Data analysis using the Langmuir 1:1 binding model (for kinetic analysis) and/or steady state analysis (for affinity analysis of data sets that reach equilibrium binding) follows data acquisition. To ensure only high-quality data are used quality control (QC) criteria (as described in the material and methods section) are applied. Examples of fitted data are shown in Figure 3, to demonstrate types of data that pass/fail the QC criteria.

**Figure 3.**
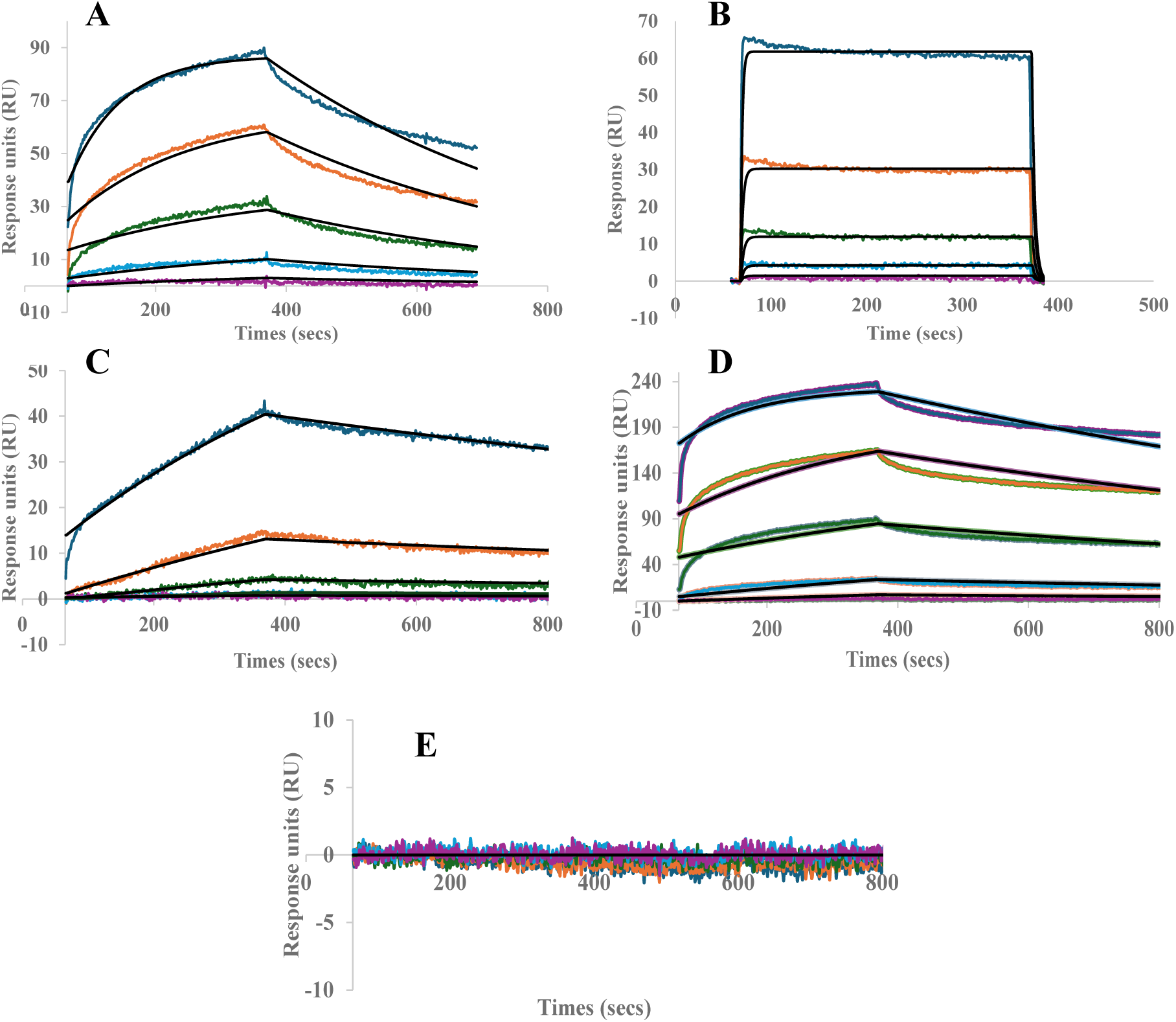
Example GAG array sensorgrams and fits. All sensorgrams are examples taken from data sets presented within this paper. Binding response from increasing concentrations of analyte are shown by coloured lines. Langmuir 1:1 fitted data are shown using black lines. **(A)** Example of kinetic data from a moderate to high affinity interaction that passes all QC checks; these data are unshaded in the GAG arrays presented in figures below. **(B)** Example of data from a low affinity interaction that passes all QC checks; these data are unshaded in the GAG arrays presented below. **(C)** Example of data where the binding response is less than 50% of R_max_ value; these data are shaded blue in the GAG arrays presented below. **(D)** Example of data where SD values are over 5% of R_max_; these data are shaded yellow in the GAG array presented below. **(E)** Example data where response is <10 RU, indicating no binding; these data are shaded grey in the GAG arrays presented below.

The known GAG-binding protein ATIII was assayed with the concentration series of 4.12–1000 nM using a 3-fold dilution series (Figure 4). Binding was observed between ATIII and heparin/HS but was not detected for CS/DS, KS or HA, which is consistent with previous findings (18,42,43). The highest affinity binding (*K*_D_) was to unfractionated heparin (UFH) and was calculated, from the experimentally determined on and off rates (*k*_a_ and *k*_d_, respectively) at approximately 96 nM (Table 3), which is consistent with previously reported data (44,45). Binding to HS was determined to be weaker with the *K*_D_ measured as 193 nM. Our analysis also suggests that an octasaccharide is close to the minimum length of heparin that is required for ATIII binding (there was no interaction with dp4), albeit that this interaction has a *K*_D_ of 575 nM. As the heparin chain length is increased the affinity between heparin and ATIII increases with the *K*_D_ of the longest oligosaccharide tested (dp24) being close to that for UFH, which is perhaps not surprising given their similar size.

The GAG array data also demonstrate that sulphation of the heparin is essential for its binding to ATIII as there was no interaction to the heparin preparation that had been fully desulphated (FdeS). The calculated *K*_D_ values for ATIII binding to all of the selectively desulphated heparins (2OdeS, 6OdeS, NdeS; Table 2) was somewhat higher than for UFH demonstrating that each sulphation group contributes to the interaction, but that neither 2-O-, 6-O- or N-sulphation is absolutely required for binding. Surprisingly, only a 2-fold difference in affinity for ATIII was seen between unfractionated heparin and HS from porcine mucosa (Table 3). In previous work using a 95-compound array of short chemoenzymatically synthesised GAGs, ATIII only bound to sequences that contained the GlcNS6S–GlcA–GlcNS3S6S–IdoA2S–GlcNS6S pentasaccharide (37), known to be required for allosteric enhancement of its enzyme activity (18,24). The HS material used here (purchased from Iduron) is described as containing both high and low sulphated species and gives a bimodal molecular weight profile with fractions of approx. 40 kDa and 9 kDa with the latter being the more sulphated component. However, while it seems unlikely that this would contain a significant amount of this rare sequence, it is apparent that this HS preparation binds to ATIII with high affinity, but where this would not be expected to cause a conformational change in ATIII, i.e., associated with increased catalytic activity.

**Figure 4.**
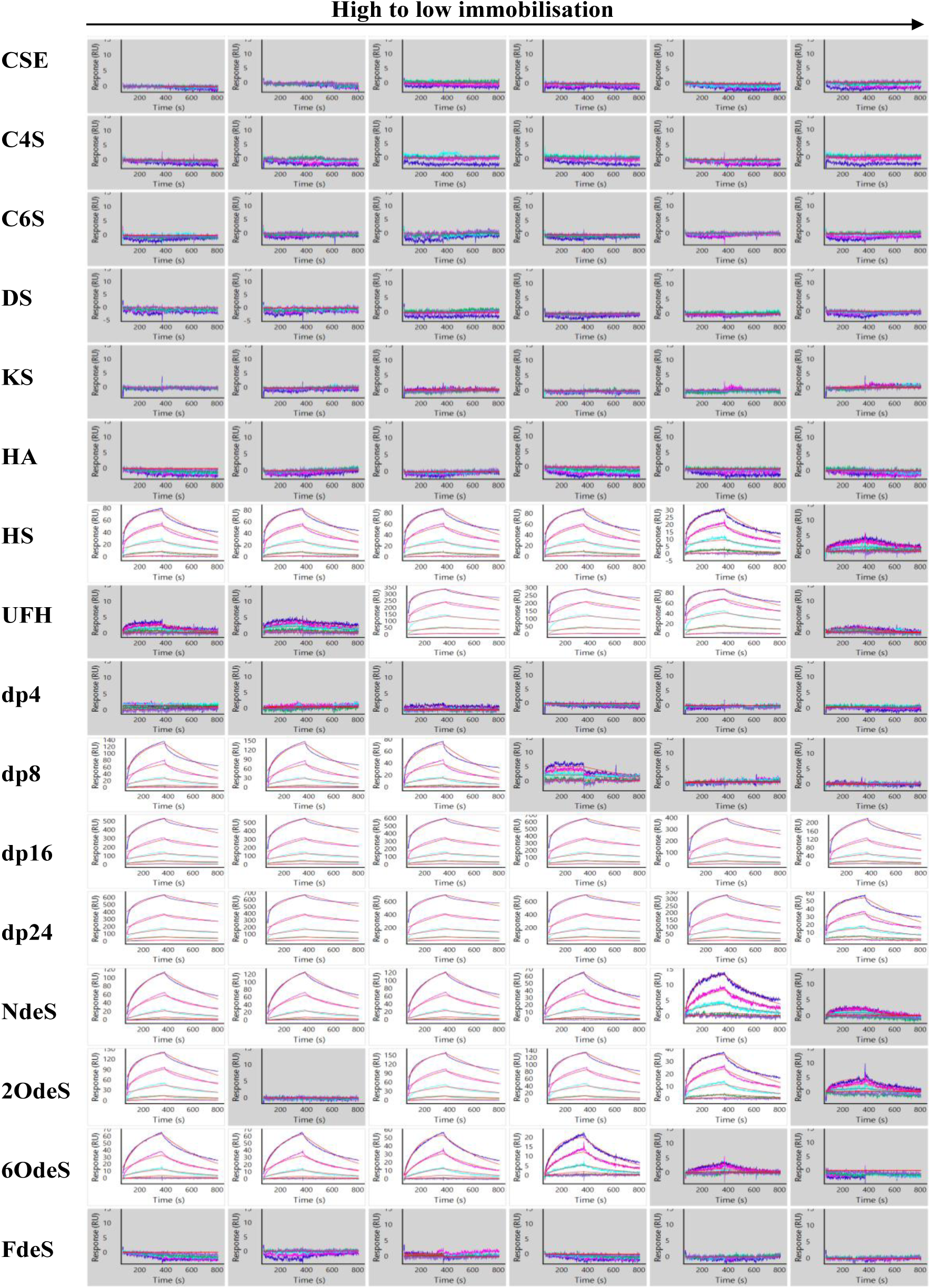
GAG array analysis of ATIII. ATIII was assayed at a concentration range from 4.12–1000 nM using a 3-fold dilution series. Binding curves are coloured as follows: 4.12 nM (yellow), 12.3 nM (purple), 37 nM (green), 111 nM (light blue), 333 nM (pink) and 1000 nM (blue). Data were analysed using the Langmuir 1:1 binding model with fitted data shown in red. GAG-protein interactions with good data fits (i.e., that pass the QC checks of analysis) are shown without shading. Grey shading denotes responses below 10 RU, indicating no binding of the protein to that GAG ‘concentration’.

**Table 3.** Parameters obtained for the ATIII-GAG interactions.

| <sup>a</sup>GAG | <sup>b</sup>n | $k_a$ (1/Ms)<sup>c</sup><br>x10<sup>4</sup> | %CV | $k_d$ (1/s)<sup>c</sup><br>x10<sup>-3</sup> | %CV | $K_D$ (M)<sup>c</sup><br>x10<sup>-7</sup> | %CV |
| --- | --- | --- | --- | --- | --- | --- | --- |
| **HS** | 5 | **1.21 ± 0.13** | 10.8 | **2.33 ± 0.23** | 9.7 | **1.93 ± 0.08** | 4.3 |
| **UFH** | 3 | **1.02 ± 0.42** | 41.3 | **0.93 ± 0.18** | 19.1 | **0.96 ± 0.18** | 18.8 |
| **dp8** | 3 | **0.49 ± 0.01** | 2.1 | **2.79 ± 0.33** | 11.9 | **5.75 ± 0.81** | 14.0 |
| **dp16** | 6 | **0.52 ± 0.01** | 1.4 | **1.19 ± 0.27** | 22.3 | **2.27 ± 0.49** | 21.4 |
| **dp24** | 6 | **0.61 ± 0.12** | 20.0 | **1.20 ± 0.69** | 57.9 | **1.87 ± 0.59** | 31.7 |
| **NdeS** | 5 | **0.53 ± 0.04** | 8.1 | **2.10 ± 0.14** | 6.7 | **3.95 ± 0.88** | 22.4 |
| **2OdeS** | 4 | **1.07 ± 0.23** | 21.9 | **1.89 ± 0.43** | 22.5 | **1.77 ± 0.04** | 2.3 |
| **6OdeS** | 4 | **1.04 ± 0.23** | 22.3 | **2.89 ± 0.32** | 10.8 | **2.94 ± 0.55** | 18.8 |

<sup>a</sup>No binding of ATIII to CSE, C4S, C6S, DS, KS, HA, dp4 and FdeS; <sup>b</sup>n = number of regions of interest (spots) from which kinetic and affinity data were derived. <sup>c</sup>Mean values ± SD.

To further test the capabilities of the array the GAG binding of inter-α-inhibitor (IαI) heavy chain 1 (HC1) was assessed. IαI is a plasma proteoglycan composed of two heavy chains (HC1, HC2) covalently linked to bikunin via a CS chain (17,46,47). At sites of inflammation (and during ovulation), HC1 (and HC2) is transferred onto HA by TSG-6 (the protein product of tumour necrosis factor-stimulated gene-6), generating covalent HC•HA complexes that crosslink HA chains (41,48–51). The non-covalent interactions of HC1 with sulphated GAGs has not been systematically assessed, although it is known to bind heparin via affinity chromatography and microtitre plate assay (29,41). Analysis here demonstrates a moderate affinity interaction of HC1 with UFH and HS of 280 nM and 440 nM, respectively (Figure 5 and Table 4). Interestingly, although HC1 can bind to a tetrasaccharide (dp4) of heparin it requires longer chains for maximal binding with the interaction being weaker (>1µM) below dp24. Selective removal of 2-O-, 6-O- or N-sulphate groups does not greatly affect the affinity of HC1 binding to heparin, however, removal of all sulphate groups (FdeS) abolished binding, showing that sulphation is required for this interaction. With the 6OdeS two different modes of binding are seen, i.e., with low and high affinity interactions seen at the lower and higher GAG-coating concentrations, respectively. The reason for this is unclear, but it might indicate that HC1 binds tightly to a rare sequence in the 6OdeS preparation.

Here screening also shows that HC1 does not interact non-covalently with HA (as was anticipated (17,50), nor to KS or DS (Figure 5). Weak binding was detected between HC1 and CSE, but not to C4S or C6S suggesting both 4-O- and 6-O sulphation are required for the interaction. However, it was not possible to obtain quantifiable data for the HC1-CSE interaction, as indicated by the blue shading in Figure 5; i.e., the concentration of HC1 used led to a response that was 50% less than the fitted R_max_, suggesting a relatively weak interaction (>2 µM); for this reason, calculated values for CSE were excluded from Table 4. The data from one concentration of UFH (shaded yellow in Figure 5), were also excluded since the fits produced residual values greater than 5% of the calculated R_max_. The remaining data for UFH did pass quality control checks, allowing quantifiable measurements that are summarised in Table 4.

**Figure 5.**
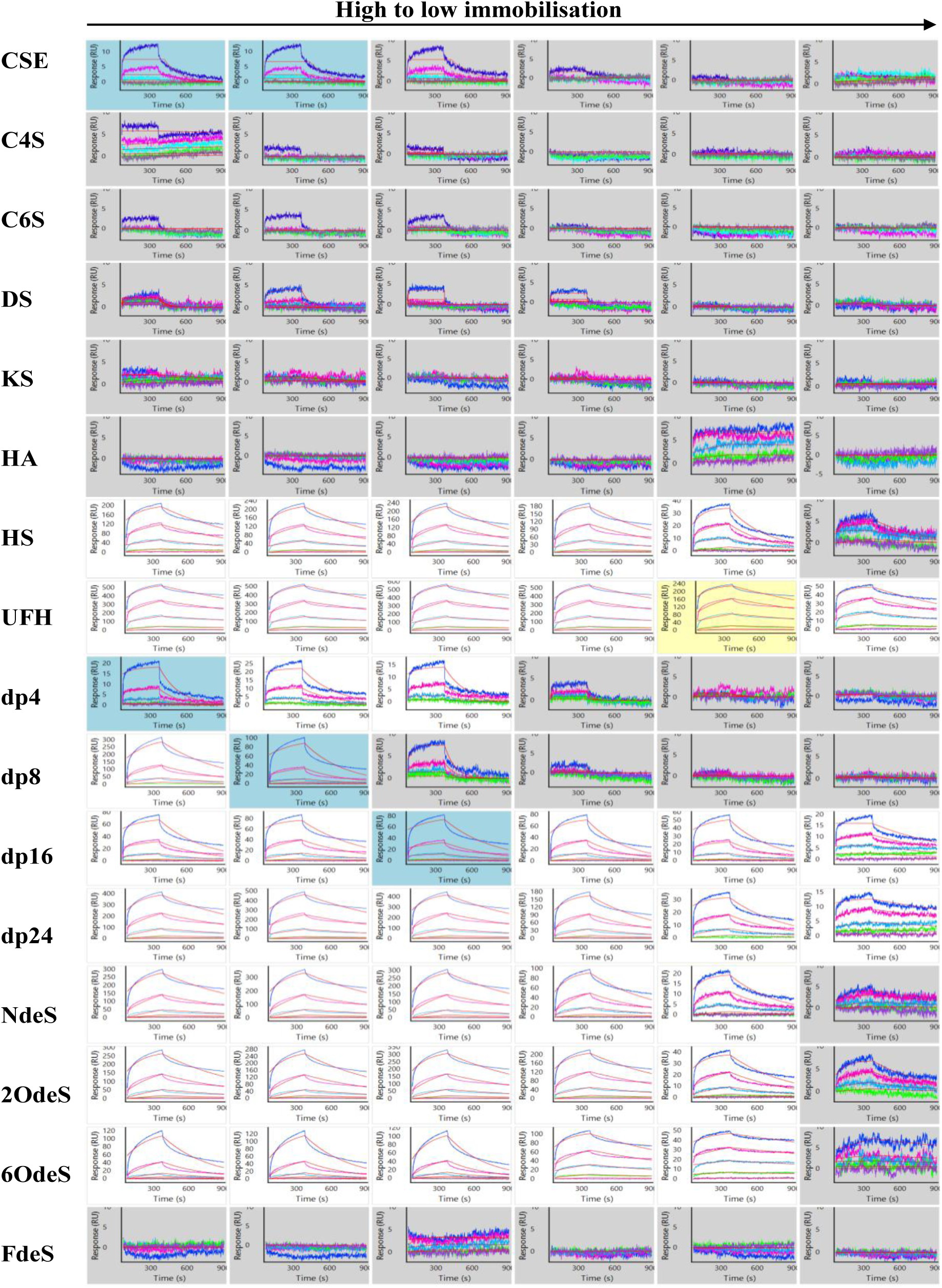
GAG array analysis of HC1. HC1 was assayed over a concentration range of 8.23–2000 nM using a 3-fold dilution series. Binding curves are coloured as follows: 8.23 nM (yellow), 24.7 nM (purple), 74.1 nM (green), 222 nM (light blue), 667 nM (pink) and 2000 nM (blue). Data were analysed using the Langmuir 1:1 binding model with fitted data shown in red. GAG-protein interactions with good data fits (i.e., that pass QC checks) are shown without shading. Grey shading denotes responses below 10 RU, indicating no binding. Blue shading denotes data that is less than 50% of R_max_ value and yellow denotes data where SD values are over 5% R_max_. Blue and yellow shaded interactions are not included in the determination of the binding parameters shown in Table 4.

**Table 4.**
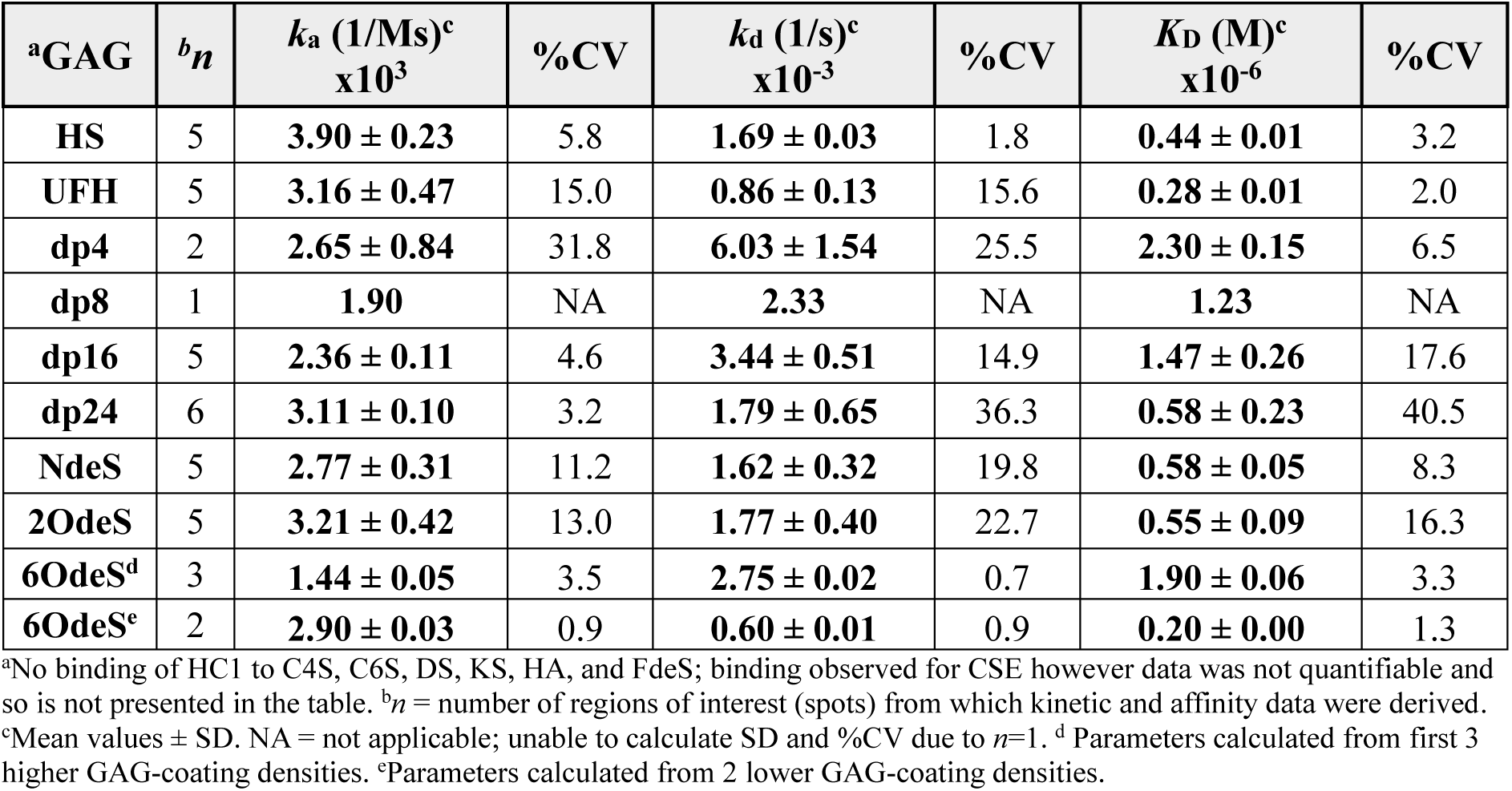
Parameters obtained for the HC1-GAG interactions.

The secreted protein Slit2 acts as a cellular guidance cue, e.g., for neurons and cells of the innate immune system, inhibiting their migration. Slit2 functions via its interaction with roundabout (robo) receptors and binds to cell-surface HS chains on HSPGs for localisation and signalling (36,52,53). Further studies have also suggested a high affinity interaction between Slit2 and KS (*K*_D_ = 28nM) (54). Our data on Slit2 reveals that the protein binds all the sulphated GAGs present within the array with no binding detected for only FdeS heparin or HA (Figure 6 and Supplementary Figure 1). Previous SPR studies (55) have determined the affinity of the Slit2-HS interaction to be either 10 nM or 50 nM for the recombinant D2 and D4 domains, respectively, where the former is close to our calculated *K*_D_ of 6 nM (Table 5). The *K*_D_ for Slit2-KS interaction was determined as 7.9 nM, which is similar to published data (54). The interaction between Slit2 and UFH was also found here to be high affinity (2.6 nM), where the selectively desulphated heparin preparations (2OdeS, 6OdeS and NdeS) bound with similar *K*_D_ values (5.7-6.7 nM), although, as noted above, FdeS, with all sulphates removed did not bind. Moreover, heparin chain length did have an effect on Slit2 binding. While the protein was able to interact with the tetrasccharide, kinetic measurements were not possible for dp4 or dp8 due to the data being <50% R_max_, indicating weak binding (Supplementary Figure 1). On the other hand, dp16 and dp24 bound to Slit2 with low nM affinities. Binding of Slit2 to CS/DS has not been reported before, but our analysis shows strong interactions with CSE (*K*_D_ = 4.9 nM) and DS (*K*_D_ =10 nM) (Table 5). Binding to CSE appears to be largely mediated by the 6-O sulphate groups, since CSE and C6S bind with similar affinities and the observed binding of Slit2 to C4S was too weak to measure (Supplementary Figure 1). Interpretation of Slit2 binding to the GAG array was impacted by levels of non specific binding to the SA reference surface. Whilst this is unlikely to have an effect on the spots with higher R_max_ values (>30RU) the need to subtract ca 30 RU of non specific binding does reduce the interpretability of the lower R_max_ interactions.

**Figure 6.**
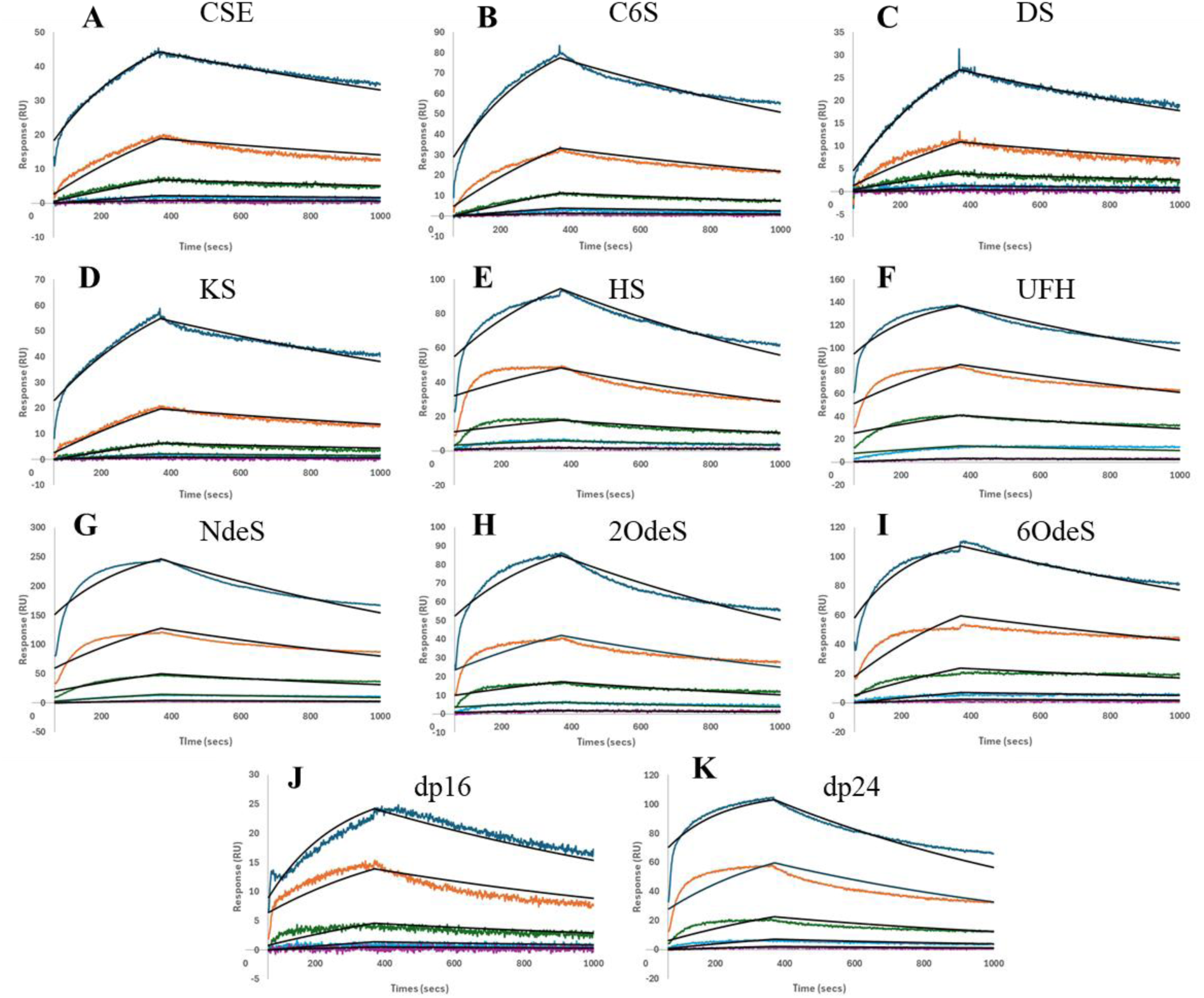
Selected sensorgrams from the GAG array analysis of Slit2. Slit2 concentration series 28 nM (dark blue), 9.3 nM (orange), 3.1 nM (green), 1 nM (blue), 0.35 nM (purple). Representative plots of Slit2 binding to CSE (A), C6S (B), DS (C), KS (D), HS (E), UFH (F), NdeS (G), 2OdeS (H), 6OdeS (I), dp16 (J), dp24 (K). Langmuir 1:1 fit shown in black.

**Table 5.** Parameters obtained for the Slit2-GAG interactions.

| <sup>a</sup>GAG | <sup>b</sup>*n* | *k<sub>a</sub>* (1/Ms)<sup>c</sup><br>x10<sup>5</sup> | %CV | *k<sub>d</sub>* (1/s)<sup>c</sup><br>x10<sup>-4</sup> | %CV | *K<sub>D</sub>* (M)<sup>c</sup><br>x10<sup>-9</sup> | %CV |
| --- | --- | --- | --- | --- | --- | --- | --- |
| CSE | 2 | 1.21 ± 0.08 | 7.0 | 5.90 ± 0.33 | 5.5 | 4.89 ± 2.19 | 44.8 |
| C6S | 2 | 1.18 ± 0.18 | 15.0 | 7.20 ± 0.61 | 8.4 | 6.25 ± 1.48 | 23.8 |
| DS | 4 | 0.88 ± 0.08 | 8.5 | 8.77 ± 0.87 | 10.0 | 10.1 ± 1.68 | 16.7 |
| KS | 1 | 0.73 | NA | 5.76 | NA | 7.90 | NA |
| HS | 4 | 1.72 ± 0.42 | 24.5 | 9.55 ± 1.03 | 10.8 | 5.96 ± 2.17 | 36.4 |
| UFH | 4 | 2.11 ± 0.32 | 15.2 | 5.40 ± 0.05 | 1.0 | 2.61 ± 0.42 | 16.1 |
| dp16 | 4 | 1.34 ± 0.13 | 9.3 | 12.3 ± 3.47 | 28.2 | 9.35 ± 3.14 | 33.6 |
| dp24 | 2 | 2.03 ± 0.04 | 1.7 | 8.62 ± 1.87 | 21.8 | 4.26 ± 0.99 | 23.2 |
| NdeS | 3 | 1.53 ± 0.35 | 22.8 | 8.44 ± 0.87 | 10.3 | 5.72 ± 1.59 | 27.9 |
| 2OdeS | 4 | 1.58 ± 0.47 | 29.8 | 8.71 ± 0.49 | 5.7 | 5.99 ± 2.16 | 36.0 |
| 6OdeS | 4 | 1.41 ± 0.33 | 23.8 | 9.01 ± 0.30 | 3.3 | 6.65 ± 1.61 | 24.2 |

<sup>a</sup>No binding of Slit2 to HA and FdeS; binding observed for C4S, dp4 and dp8 however data was not quantifiable and so are not presented in the table (see Supplementary Figure S1). <sup>b</sup>*n* = number of regions of interest (spots) from which kinetic and affinity data were derived. <sup>c</sup>Mean values ± SD. NA = not applicable; unable to calculate SD and %CV due to *n*=1.

CD44 is a ubiquitous multifunctional transmembrane glycoprotein that acts as a primary cell-surface receptor for HA supporting numerous cellular processes including adhesion, migration, proliferation and survival (17,56,57). The CD44-HA interaction has been previously characterised as being in the low micromolar range (34,58,59). In addition, CD44 has also been reported to bind to C4S and C6S (57,60,61), however, no competition of HA binding was seen with ultrapure preparations of C4S or C6S (59), suggesting that if these interactions occur they are not mediated by the same site. Here using our recombinant HA-binding domain from human CD44 (CD44_HABD^20-169^) (16) we measured its affinity for HA12 (Figure 7, Supplementary Figure 2 and Table 6). As equilibrium binding was reached at each concentration, we were able to perform both kinetic and steady state analysis. The affinity was calculated to be around 30 µM, as determined by both kinetic and steady state analysis; a comparable value of ∼60 µM was obtained by isothermal titration calorimetry for a related construct (CD44_HABD^20-178^) with HA10 (34). The GAG array analysis showed that CD44 does interact with C4S (∼25 µM) and CSE (∼29 µM) and with similar affinities to HA; C6S was also concluded to bind, however, this interaction (that is likely in the same order of magnitude) was not quantifiable, but can be safely assumed to be > 30 uM. (Supplementary Figure 2). CD44 did not bind to any of the other GAG preparations, revealing that this receptor is specific for HA and CS.

**Figure 7.**
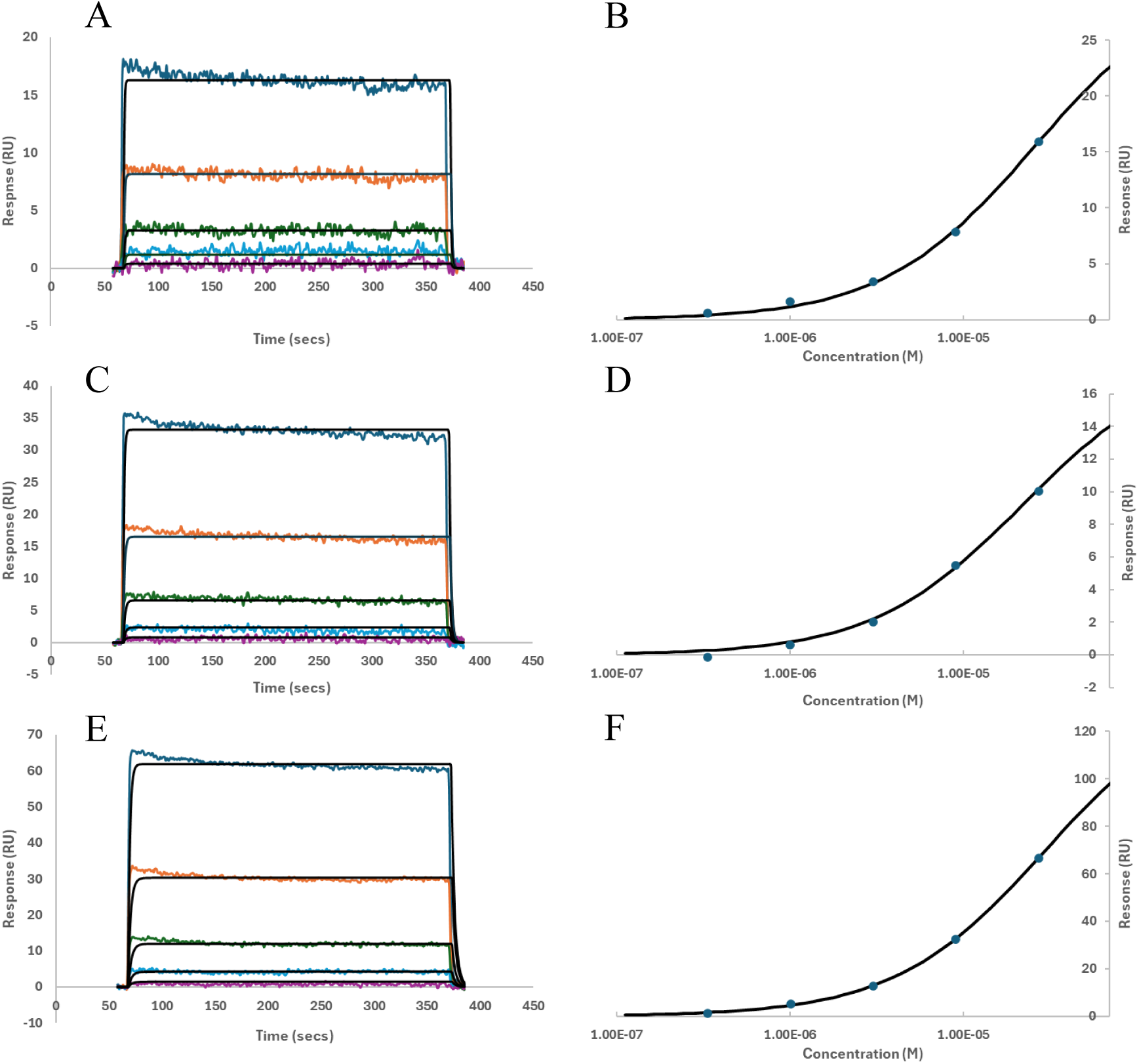
Selected sensorgrams from the GAG array analysis of CD44. CD44 was assayed over a concentration range of 0.336-27.3 µM using a 3-fold dilution series. Representative binding curves are coloured as follows: 0.336 µM (purple), 1.01 µM (light blue), 3.03 µM (green), 9.08 µM (orange) and 27.3 µM (blue). Data were analysed either using the 1:1 binding model (A, C, E) or via steady state non-linear regression (B, D, F) for CSE (A, B), C4S (C, D) and HA (E, F). Fitted data are shown in black.

**Table 6.**
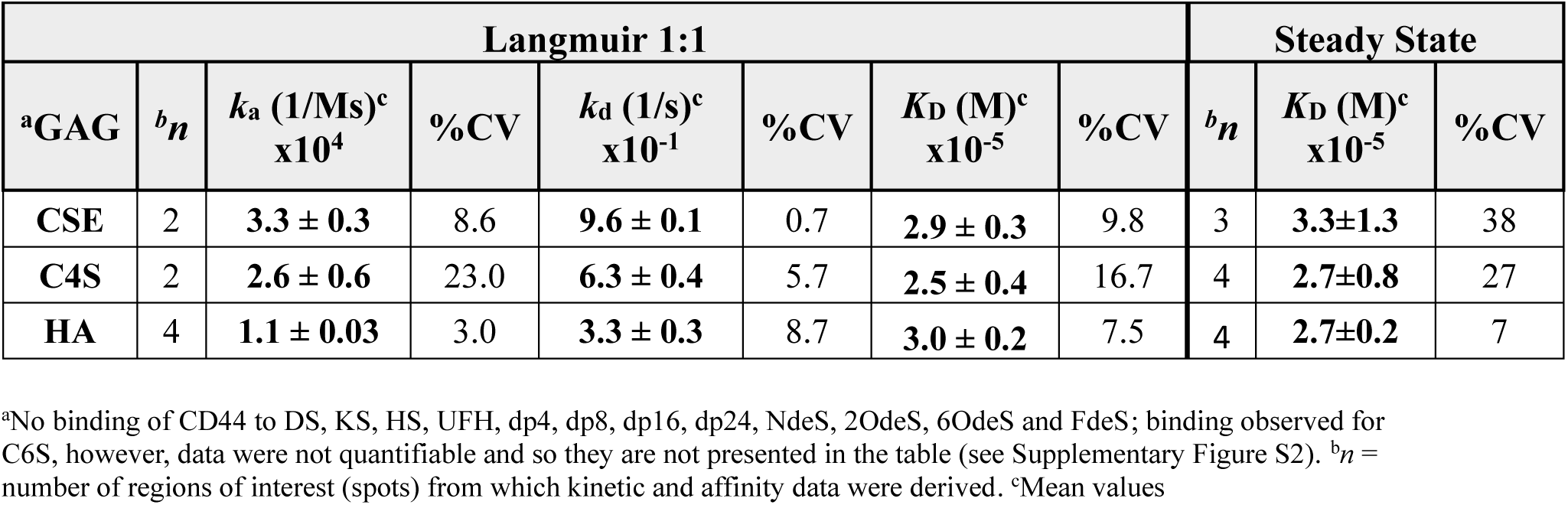
Parameters obtained for the CD44-GAG interactions.

## Discussion

Protein-GAG interactions are essential to numerous biological processes, and it has been estimated that there are >3,500 unique GAG-binding proteins encoded in the human genome (9,10). While there is a wealth of data on particular GAG-protein interactions, there is currently limited ways to systematically study their specificities and only laborious methods to generate fully quantitative data. In recent years, there has been much progress in the development of microarray formats utilising immobilised GAGs (25) where these can provide insights into specificity, and for example the use of fluorescence intensity to estimate *K*_D_ values (37).

Within this paper we have presented a novel high throughput SPR-based technique for the multiplex analysis of GAG-protein interactions. We have demonstrated the use of this GAG array to screen protein binding to multiple GAGs, identifying GAG-binding specificities, whilst simultaneously measuring the kinetics of the interactions (i.e., on and off rates) allowing the calculation of affinities (*K*_D_ values).

The technique has been validated using proteins with previously determined GAG binding specificities: ATIII (heparin), HC1 (heparin/HS), Slit2 (heparin/HS and KS) and CD44 (HA and CS). Further to this the technique was able to quantify these interactions, and the affinities obtained were largely in line with previously obtained values. Some GAG-binding specificities that had been identified in alternative techniques were however not identified, namely Slit2’s preference for HS sulphation sequences that contain 6-O-sulphation and N-sulphation (36); we observed only a 2.5- and 2.2-fold reduction in affinity with heparin that had 6-O- and N-sulphation selectively removed, and a similar 2.3-fold reduction on removal of the 2-O-sulphates. The reason for these discrepancies is unknown but could be due to differences in experimental set up such as the presentation of GAGs. In the microarray format used in (36) GAGs were noncovalently immobilised on poly-L-lysine coated slides, whereas in our array GAGs were attached to streptavidin-coated surfaces via a biotin moiety at their reducing end, meaning all of the sulphate groups are accessible rather than some being involved in ionic interactions with the surface.

Further to validating the specificity of previously known binding partners, new unknown GAG binding capabilities were identified for HC1 (CSE), Slit2 (CS/DS) and CD44 (CSE). As well as identifying these novel binders the array was able to simultaneously calculate the affinity for these interactions. The ability to identify and reliably quantify novel interactions within a single experiment is a major advantage to this technique. As noted above, other methodologies have been developed that are able to perform high throughput screening but unable to fully quantify the interaction, namely microarrays and proteomics (25,37,62–64). Conversely, information rich techniques such as NMR or traditional SPR have very low throughput and are time consuming. The GAG array presented here combines both an information rich and high throughput technique.

The throughput of our GAG array is of course substantially lower than microarrays and proteomics. However, the time saving obtained through not having to further validate and quantify binding ‘hits’ is advantageous. Furthermore, we have standardised a GAG interaction system that will allow direct comparisons across multiple research projects through use of a single defined technique with the same GAGs. In the current format, we can analyse the binding of a target protein to 16 different GAG preparations where at least 4 different proteins can be analysed on each chip. The 384-spot array could be used differently, e.g., with the analysis of one protein binding to 64 different GAG preparations, while retaining each GAG at 6 different coating concentrations. Moreover, we have only used purified GAGs (that are mostly commercially available), which in some cases have been fragmented or treated chemically to remove sulphates. Going forward we could utilise chemoenzymatically synthesised GAGs (33,65–67) in order to create bespoke arrays that allow a focus on particular research questions.

There are limitations to the technique, a number of which are inherent to the study of GAG-protein interactions by SPR (31). Some of the identified interactions failed quality control checks with regard to high standard deviation of the model fit, indicating poorly fitting data and evidence of heterogenous binding profiles is common within the analysis. Most GAGs are, as previously discussed, very heterogeneous molecules, e.g., with varied sulphation patterns, such that multiple interactions contribute to the overall binding affinity. Therefore, a high proportion GAG-protein interactions are unlikely to strictly adhere to a 1:1 binding model. Currently the Langmuir 1:1 binding model was applied to best describe the totality of the interaction, as whilst complex models with multiple on and off rates would have certainly allowed for better fitting, the results might be less meaningful to interpret as there would be no way to link the various rate constants to specific discreet interactions. The use of ‘single sequence’ chemoenzymatically synthesised GAGs may significantly reduce the kinetic complexity of these measurements.

This GAG array technique was also limited in its ability to measure weaker affinities as demonstrated by the CD44-CS interactions. This will be in part due to the concentration series used for analysis but is also an intrinsic limitation of SPR in general. That said, for the four proteins analysed here the array generated data from interactions with low nM to µM affinities, and it would be anticipated that higher affinity interactions (e.g., >50 pM) could be readily measured.

As the number of GAG-related studies increases, new techniques are required to push the field forward. The array presented here will help to expand knowledge regarding existing and newly identified GAG-protein interactions to ultimately help extend and decipher the GAG interactome. The array is available for use by the proteoglycan community, and we encourage any interested researchers to get in contact to discuss access.

## Acknowledgements

We would like to acknowledge the funding of the core facility by the Manchester Cell-Matrix Centre, particularly the Wellcome Trust Discovery Research Platform for Cell Matrix Biology (226804/Z/22/Z). AJD also acknowledges support from Arthritis UK (programme grant 22277) and the ‘Glycoweb’ BBSRC sLoLa (BB/Y00311X/1).

