## Supplemental figures for "A High-Throughput SPR-Based Array for Quantitative Profiling of Glycosaminoglycan–Protein Interactions"

Supplementary

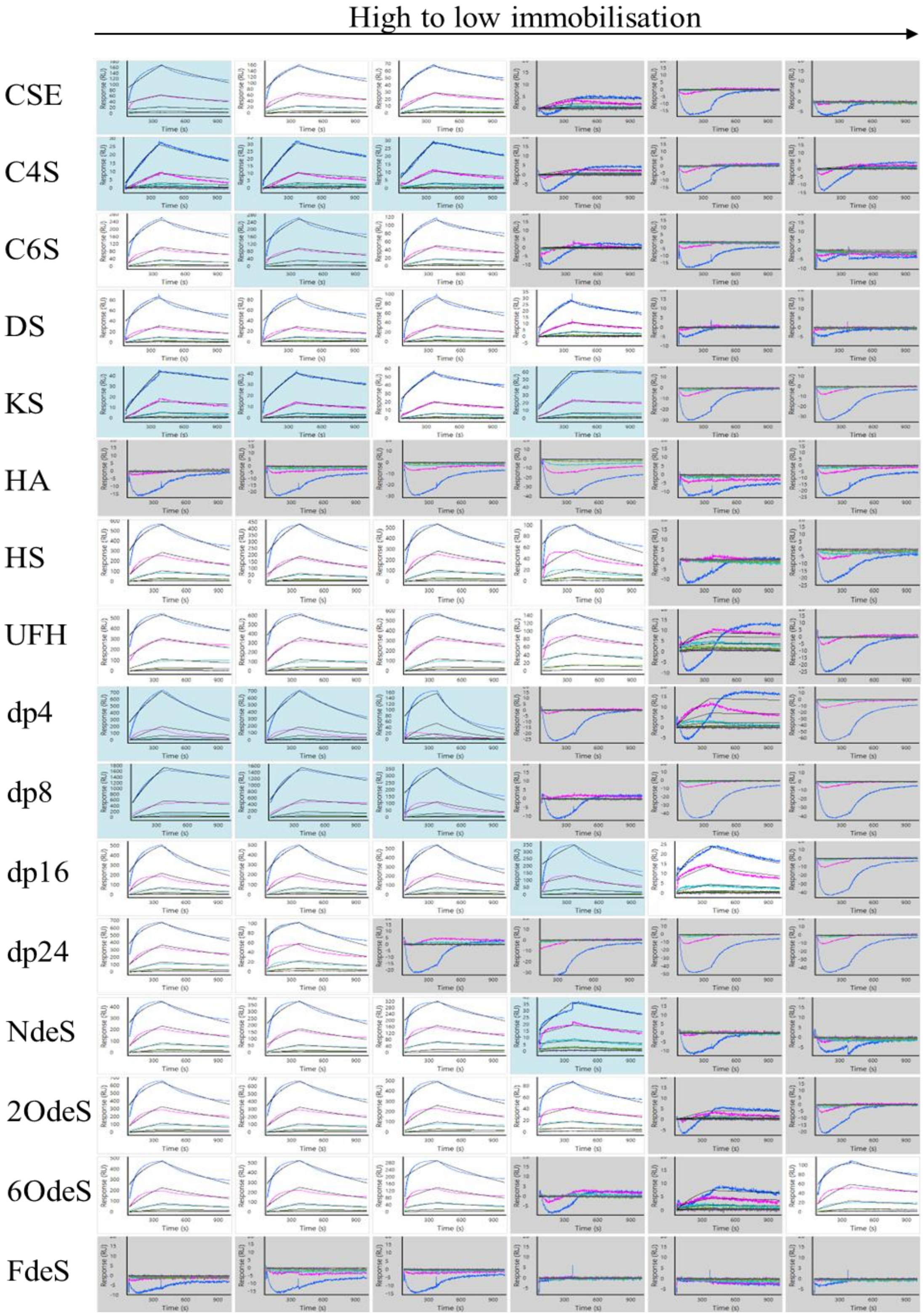


**Supplementary Figure 1. GAG array analysis of Slit2.** Slit2 assayed at a concentration series of 0.114 nM – 27.8 nM using a 3-fold dilution. Binding curves coloured as follows; 0.114 nM (yellow), 0.343 nM (purple), 1.03 nM (green), 3.09 nM (light blue), 9.26 nM (pink), 27.8 nM (blue). Data analysed using Langmuir 1:1 binding model. Fitted data shown in black. Grey shading denotes responses below 10 RU, indicating no binding. Blue shading denotes data that is less than 50% of $R_{max}$ value and are not included in determination of the binding parameters shown in Table 5.

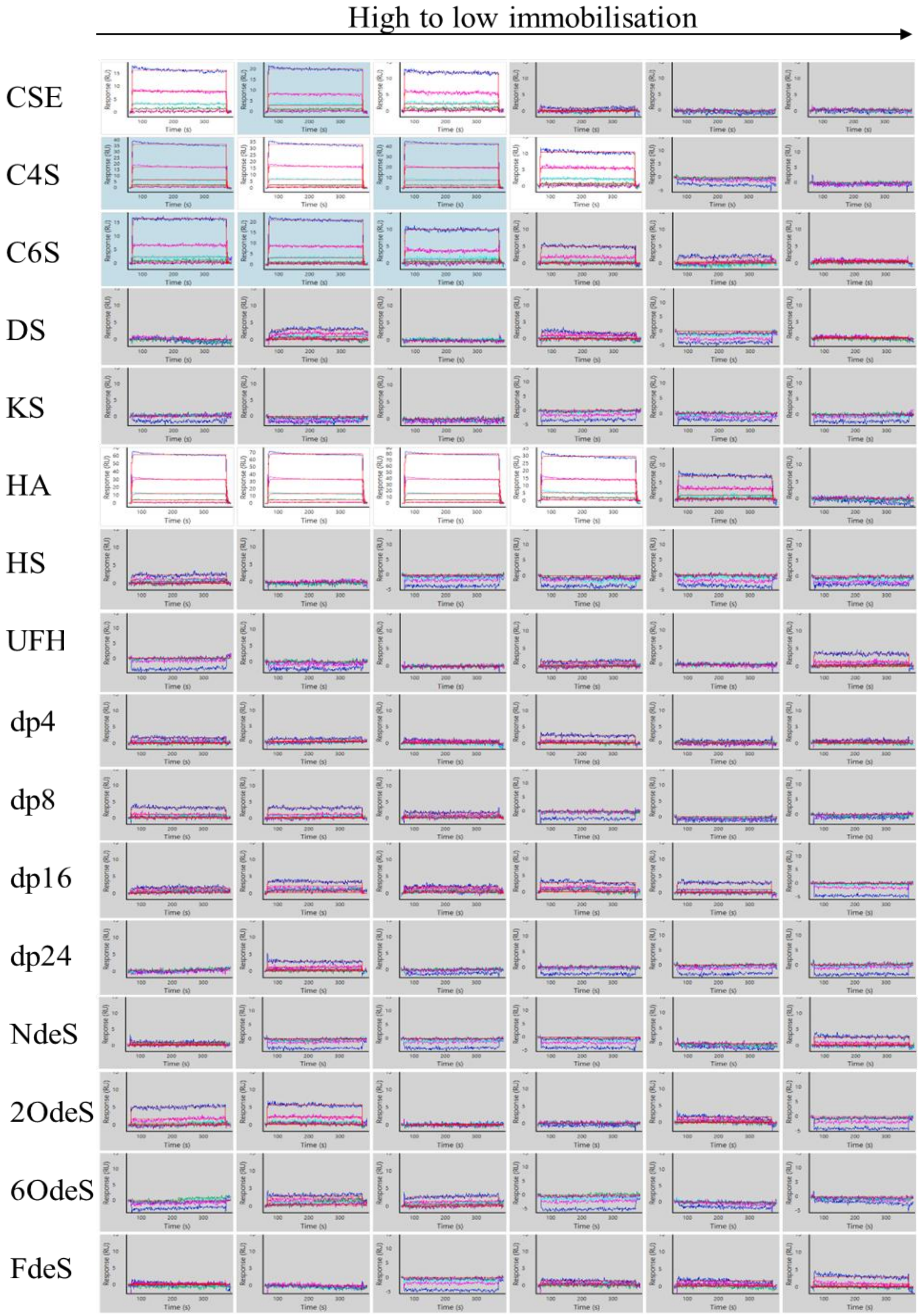


**Supplementary Figure 2. GAG array analysis of CD44.** CD44 assayed at a concentration series of 0.336 µM – 27.3 µM using a 3-fold dilution. Binding curves coloured as follows; 0.336 µM (purple), 1.01 µM (green), 3.03 µM (light blue), 9.08 µM (pink), 27.3 µM (blue). Data analysed here using Langmuir 1:1 binding model. Fitted data shown in red. Grey shading denotes responses below 10 RU, indicating no binding. Blue shading denotes data that is less than 50% of $R_{max}$ value and yellow denotes data where SD are over 5% $R_{max}$ and are not included in determination of the binding parameters shown in Table 4.
